# Neutrophil motility is regulated by both cell intrinsic and endothelial cell ARPC1B

**DOI:** 10.1101/2023.11.16.567429

**Authors:** Ashley Peterson, David Bennin, Michael Lasarev, Julia Chini, David J. Beebe, Anna Huttenlocher

**Affiliations:** Department of Medical Microbiology and Immunology, University of Wisconsin-Madison, Madison, WI, USA; Comparative Biomedical Sciences Graduate Program, University of Wisconsin-Madison, Madison, WI, USA; Department of Pediatrics, University of Wisconsin-Madison, Madison, WI, USA; Department of Biostatistics and Medical Informatics, University of Wisconsin-Madison School of Medicine and Public Health, Madison, WI, USA; Department of Pediatrics, Perelman School of Medicine, University of Pennsylvania, PA, USA; Department of Biomedical Engineering, University of Wisconsin-Madison, Madison, WI, USA; Department of Pathology and Laboratory Medicine, University of Wisconsin-Madison, Madison, WI, USA; Carbone Cancer Center, University of Wisconsin-Madison, Madison, WI, USA

**Author notes:** Corresponding author (AH).

**Keywords:** migration, neutrophil, endothelial cells, Arp2/3, PLB-985 cells, iPSC-derived neutrophils

## Abstract

Neutrophil directed motility is necessary for host defense, but its dysregulation can also cause collateral tissue damage. Actinopathies are monogenic disorders that affect the actin cytoskeleton and lead to immune dysregulation. Deficiency in ARPC1B, a component of the ARP2/3 complex, results in vascular neutrophilic inflammation; however, the mechanism remains unclear. Here we generated ARPC1B-deficient human iPSC-derived iNeutrophils that show impaired migration and a switch from pseudopodia to the formation of elongated filopodia. We show, using a blood vessel on a chip model, that primary human neutrophils have impaired movement across an endothelium deficient in APRC1B. We also show that the combined deficiency of ARPC1B in iNeutrophils and endothelium results in further reduction in neutrophil migration. Taken together, these results suggest that ARPC1B in endothelium is sufficient to drive neutrophil behavior.

**Summary statement:** The actin regulator ARPC1B in both neutrophils and endothelium drives neutrophil motility.

## Introduction

Neutrophils mediate host defense and are the first responders to infection or tissue injury. Rapid recruitment of neutrophils requires the dynamic turnover of actin at the leading edge to mediate polarized pseudopod formation and directed migration (Fritz-Laylin et al., 2017; Lämmermann et al., 2008). Defects in signaling pathways that mediate neutrophil motility result in the development of primary immune deficiencies including leukocyte adhesion deficiency (LAD)s (Dupré & Praunier, 2023). Indeed, a group of disorders characterized by abnormal actin cytoskeletal regulation have been referred to as actinopathies and lead to immune dysregulation (Papa et al., 2021). This includes defects in actin-binding proteins such as Wiskott Aldrich syndrome protein (WASP), which plays a critical role in nucleating actin filaments; its deficiency leads to both immune deficiency and autoimmunity (Ngoenkam et al., 2021; Thrasher & Burns, 2010; Zhang et al., 2006). The susceptibility to infection in patients with these disorders is partially mediated by altered leukocyte motility. However, how defects in neutrophil motility contribute to autoimmunity and vascular inflammation remains less clear.

Neutrophil chemotactic migration involves the asymmetric distribution of dynamic F-actin at the leading edge that is regulated by a complex of actin regulatory proteins including WASP. WASP promotes actin filament branching at the cell front through its association with the actin-related protein 2/3 complex (Arp2/3). The Arp2/3 complex is comprised of 7 components including the actin regulatory protein ARPC1B, that binds directly to WASP (Leung et al., 2021) . Deficiency in ARPC1B is associated with immune dysregulation in children characterized by recurrent infections, vasculitis and allergies. There have been recent studies focused on the role of ARPC1B in B-cell function (Leung et al., 2021), T-cell function (Brigida et al., 2018), and a recent report showed that there is a defect in ARPC1B patient neutrophil transendothelial migration (Kempers et al., 2021). While this recent study examined the effects of ARPC1B deficiency on neutrophil transendothelial migration, the authors used only patient neutrophils in combination with unmodified endothelial cells. To date, the role of ARPC1B in endothelial cells remains unknown. However, one key feature of ARPC1b deficiency is leukocytoclastic vasculitis (Kahr et al., 2017; Kuijpers et al., 2017). This syndrome is characterized by a dense neutrophil infiltrate in and around blood vessels characteristic of endothelial inflammation. It is unclear what drives the vasculitis and if it is mediated specifically by a neutrophil-intrinsic defect in ARPC1B expression. To address this gap, we aimed to use microfluidic systems to dissect the function of ARPC1B in both neutrophils and endothelial cells. We compared the effects of ARPC1B deficiency using both the established promyelocytic leukemia cell lines, PLB-985 (Collins et al., 1977; Saito et al., 1978), and a more recent human iPSC-derived neutrophil model that also enables the study of ARPC1B deficient endothelium (Brok-Volchanskaya et al., 2019).

Here we examined the role of ARPC1B in both neutrophils and endothelial cells during dynamic neutrophil-endothelial interactions. We used the human iPSC system to model ARPC1B deficiency. We found that ARPC1B-deficient iPSC-derived neutrophils (iNeutrophils) more closely recapitulate primary patient neutrophils in terms of the motility defect and actin cytoskeletal changes. In addition, we found that endothelial cell ARPC1B plays a key role in regulating neutrophil dynamic interactions with the endothelium.

## Results

### ARPC1B-deficient PLB-985 neutrophil-like cells have a modest motility defect

Previous studies have shown that neutrophils from patients with ARPC1B deficiency display altered neutrophil motility and organization of the actin cytoskeleton (Kempers et al., 2021; Kuijpers et al., 2017). To further characterize the role of ARPC1B in neutrophil function, we used the myeloid leukemia cell line, PLB-985 cells, which can be terminally differentiated into neutrophil-like cells using DMSO (Collins et al., 1978). Using CRISPR mutagenesis, we generated ARPC1B knock out PLB-985 cells that were confirmed by western immunoblotting (Fig. 1A,B). We assessed the motility of these cells using previously described microfluidics devices (Berthier et al., 2010; Yamahashi et al., 2015) and live cell imaging. Consistent with ARPC1B-deficient patient neutrophils, ARCP1B-deficient PLB-985 cells had reduced cell speed, although there were no statistically significant changes in track length or persistence (Fig. 1C, D). To determine the mechanism of this defect we examined the organization of the actin cytoskeleton in the knockout cells. The ARPC1B KO cells had reduced integrated density of F-actin in response to fMLP (Fig. 1E,F). However, unlike patient neutrophils there was only a modest increase in the number of filopodia per cell and no change in filopodia length (Fig. 1F-H). Taken together, PLB-985 cells deficient in ARPC1B display impaired migration and exhibit actin cytoskeletal changes consistent with the defect reported in neutrophils from patients deficient in ARPC1B. However, the phenotypes were only modest, suggesting that this cancer cell line has some compensatory mechanisms that enable motility and actin cytoskeletal reorganization in the absence of ARPC1B.

**Figure 1.**
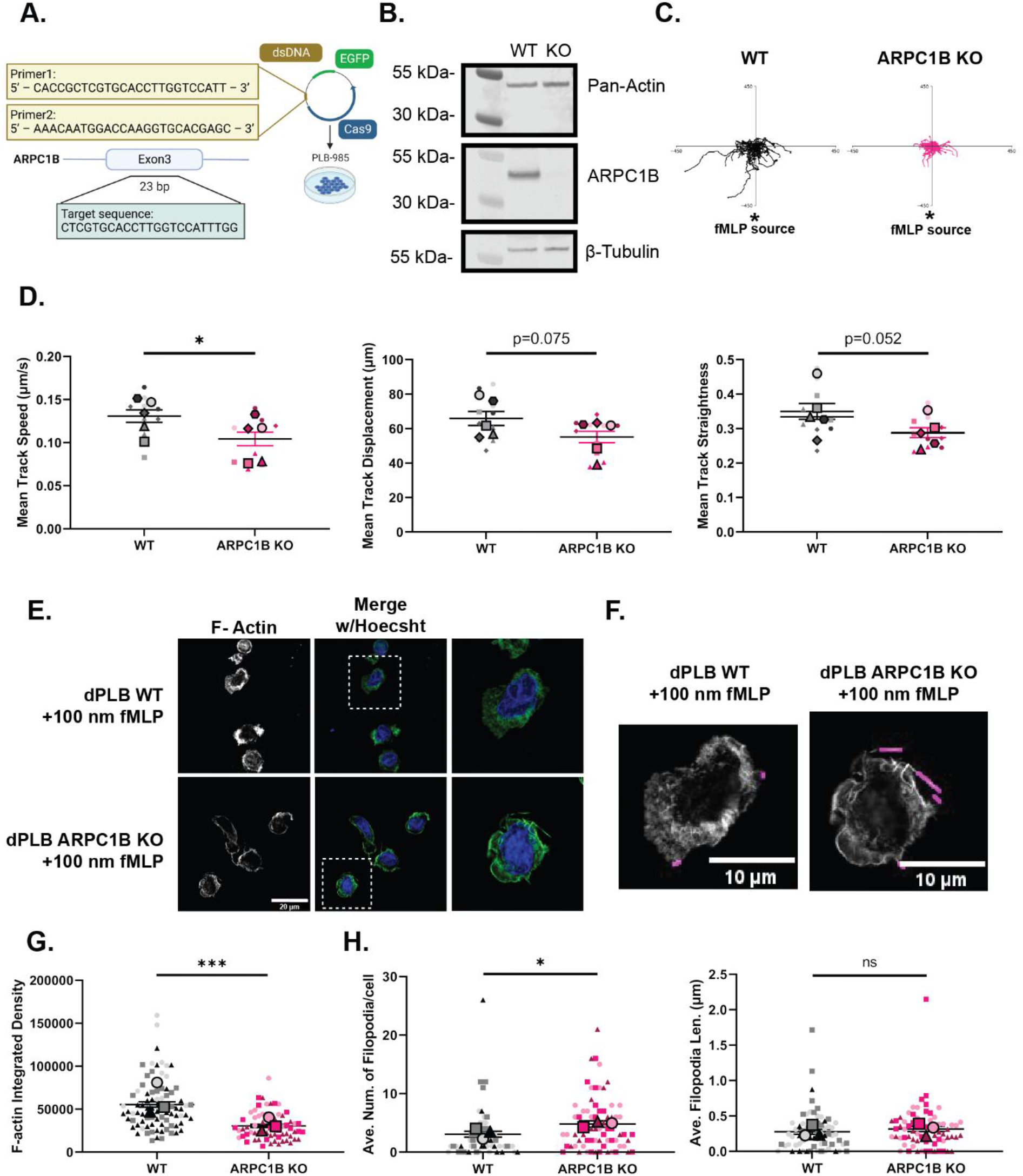
ARPC1B-deficient PLB-985 cells display a defect in directed migration. (A) Diagram illustrating sgRNAs targeting exon 9 of ARPC1B in undifferentiated PLB-985 cells. Diagram created in BioRender. (B) Western blot to confirm deletion of ARPC1B in differentiated PLB-985 cells. (C) Representative rose plots showing motility of ARPC1B-/-differentiated PLB-985 cells compared to WT. Plots are oriented with an fMLP gradient originating at the bottom. (D) Motility of differentiated PLB-985 cells as quantified by mean track speed, mean track displacement, and mean track straightness. Track metrics are averaged across all tracks in a microfluidics device, and then further averaged within each day. (E) Representative immunofluorescence staining of F-actin and Hoechst nuclear stain with 100 nm uniform fMLP stimulation for 15 minutes. (F) Representative filopodial detection using FiloQuant ImageJ plugin. (G) Quantification of F-actin integrated density. Plots in (G) and (H) represent biological replicate averages superimposed on individual cell quantifications for each day. Means ± SEM are shown. p values were calculated by Student’s t-test (D) or linear mixed effects model (G) and (H) *, p < 0.05; **, p < 0.01; ***, p < 0.001; ****, p < 0.0001.

### ARPC1B-deficient IPSC-derived neutrophils have increased filopodia formation and impaired migration

To determine if the patient phenotype could be recapitulated using a human iPSC system, we depleted ARPC1B from human pluripotent stem cells (iPSC) using CRISPR mutagenesis (Fig. 2A, B). In prior studies we have shown that GMP compatible iPSC-derived neutrophils, referred to as iNeutrophils, have motile function and can be genetically engineered to display improved migration in response to gradients of chemoattractant (Brok-Volchanskaya et al., 2019a; Giese et al., 2023). ARPC1B KO and WT iNeutrophils had similar expression of neutrophil maturity markers such as CD11b, CD15 and CD16, and other surface makers (Sup. Fig. 1A,B). We also generated hemogenic endothelial cells during an early step in the hematopoietic differentiation protocol and confirmed depletion of ARPC1B in the hemogenic endothelium (Fig. 2B). We found that ARPC1B-deficient iNeutrophils displayed a significant defect in motility with reduced speed, track displacement and length compared to WT iNeutrophils (Fig. 2C, D). Similar to patient neutrophils (Kahr et al., 2017; Kuijpers et al., 2017; Volpi et al., 2019), ARPC1B-deficient iNeutrophils showed a loss of cell polarity in response to fMLP with the formation of long filopodia (Fig. 2E, F). Quantification showed a significant increase in the length and number of filopodia in the ARPC1B-deficient cells (Fig. 2G,H). Taken together these findings suggest that the iPSC-derived neutrophils can be used to model ARPC1B deficiency and more closely recapitulate the human disease phenotype than PLB-985 cells.

**Figure 2.**
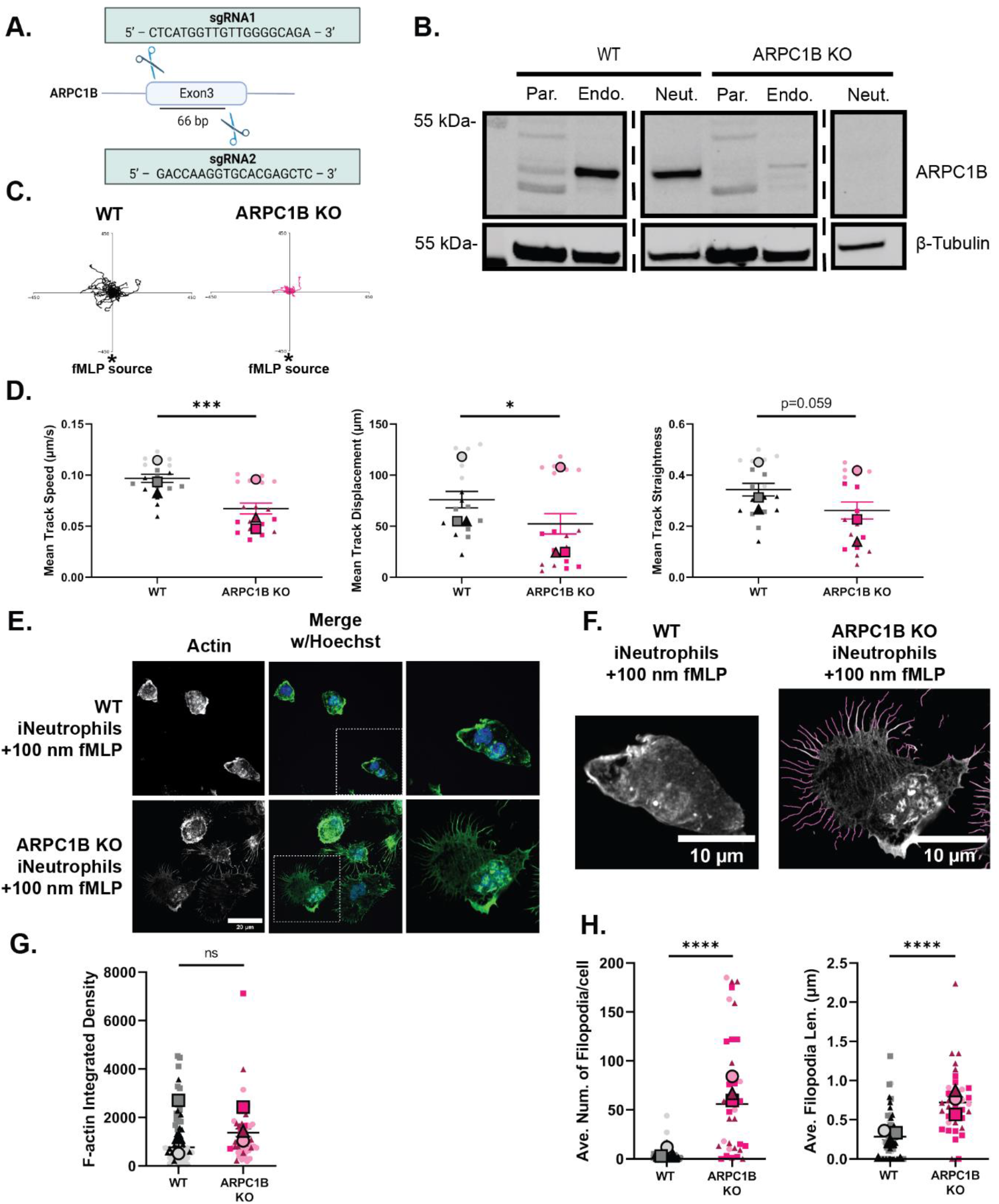
ARPC1B-deficient iNeutrophils have impaired directed migration and display a switch from pseudopodia to filopodia. (A) Diagram illustrating sgRNAs targeting exon 9 of ARPC1B at the pluripotent stem cell stage. Diagram created in BioRender. (B) Western blot to confirm deletion of ARPC1B in differentiated neutrophils and endothelial cells. (C) Representative rose plots showing motility of ARPC1B-/-iNeutrophils compared to WT. Plots are oriented with an fMLP gradient originating at the bottom. (D) iNeutrophil motility as quantified by mean track speed, mean track displacement, and mean track straightness. Track metrics are averaged across all tracks in a microfluidics device, and then further averaged within each day. (E) Representative immunofluorescence staining of F-actin and Hoechst nuclear stain with 100 nm uniform fMLP stimulation for 15 minutes. (F) Representative filopodial detection using FiloQuant ImageJ plugin. (G) Quantification of F-actin integrated density. Plots in (G) and (H) represent biological replicate averages superimposed on individual cell quantifications for each day. Means ± SEM are shown. p values were calculated by Student’s t-test (D) or linear mixed effects model (G) and (H) *, p < 0.05; **, p < 0.01; ***, p < 0.001; ****, p < 0.0001.

### ARPC1B-deficient IPSC-derived endothelial cells regulate neutrophil behavior

A hallmark of ARPC1B deficiency is the presence of vasculitis. The mechanism of the vasculitis remains largely unknown. However, previous studies have suggested a defect in the transendothelial migration of ARPC1B-deficient patient neutrophils (Kempers et al., 2021). Here we sought to use the iPSC system to determine if endothelial cell ARPC1B modulates neutrophil behavior. ARPC1B is expressed in iPSC-derived endothelial cells (iECs) (Fig. 2B), however its function remains unknown as previous work has focused on motile cells of the immune system. We harvested the endothelial cells generated early in the iNeutrophil differentiation process and verified that there were no significant difference in endothelial maturity and identity between WT and ARPC1B KO endothelium (Sup. Fig.1B). To address the role of ARPC1B in endothelial cell function we used our previously described LumeNext blood vessel on a chip model (Ingram et al., 2018). We seeded either WT or ARPC1B KO iECs to form a blood vessel lumen and then loaded the lumens with primary human neutrophils from healthy donors to quantify movement to a source of *Pseudomonas Aeruginosa,* using published methods (Hind et al., 2018; Ingram et al., 2018)(Fig. 3A). In the lumen model the ARPC1B-deficient endothelium formed a lumen with similar permeability to control endothelium when exposed to *P. Aeruginosa* (Fig 3B), suggesting that the endothelium remains intact in the absence of ARPC1B. We found that knockout of ARPC1B in endothelial cells impaired the transmigration and migration of primary blood neutrophils out of the vessel toward *P. Aeruginosa* (Fig. 3C). The neutrophils remained associated with the endothelium, a hallmark of neutrophil vasculitis. We also found that ARPC1B-deficient endothelium generates more cytokines including IL-1β and TNF-α on 2D surface (Fig 3E), which indicates a more inflammatory endothelium. Taken together, these findings suggest that ARPC1B in endothelial cells impairs neutrophil transmigration in the LumeNext blood vessel on a chip. This is, to our knowledge, the first evidence that ARPC1B in endothelium regulates primary human neutrophil behavior.

**Figure 3.**
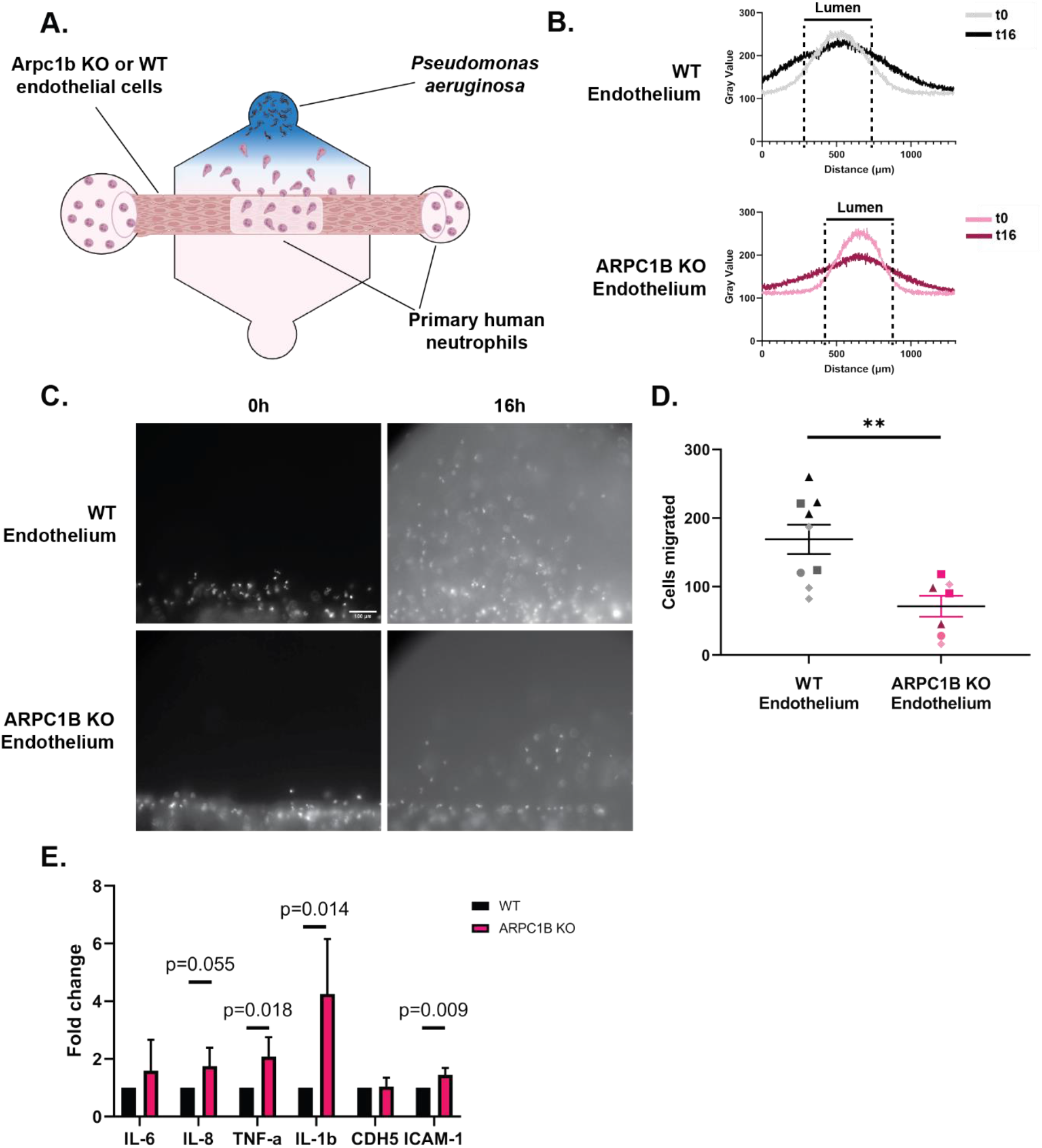
ARPC1B-deficient endothelial cells modulate primary human neutrophil motility and have increased inflammatory gene expression. (A) Schematic showing the setup of LumeNext devices with WT or ARPC1B -/-endothelial cells. (B) Endothelial permeability of lumens seeded with WT or ARPC1B-/-cells. (C) Representative images of fluorescently labeled primary neutrophil motility at 0 hours and 16 hours out of endothelial lumens. (D) Quantification of the number of cells that migrate out of the lumens after 16 hours. (E) qPCR analysis of inflammatory gene transcription. p values were calculated by Student’s t-test *, p < 0.05; **, p < 0.01; ***, p < 0.001; ****, p < 0.0001.

### Both ARPC1B in iNeutrophils and endothelium modulate neutrophil 2D migration

To determine if both ARPC1B in endothelium and iNeutrophils affect neutrophil motility, we used a 2-dimensional system to image the migration of iNeutrophils on endothelium. This was particularly important because WT iNeutrophils and PLB-985 cells are not able to migrate out of the endothelium in the LumeNext model (data not shown). To image neutrophil motility on endothelium, we seeded WT or ARPC1B KO iNeutrophils on confluent monolayers of WT or KO endothelium and found that the phenotype of the neutrophils was different depending on the combination of KO cells. We found more aggregation of iNeutrophils on the combined KO background compared to the single KO in either iNeutrophils or endothelium (Fig. 4A). Accordingly, we found that the WT iNeutrophils were highly motile on WT endothelium and migration was impaired with the ARPC1B-deficient iNeutrophils (Fig 4B) with reduced mean track velocity and distance migrated (Fig 4C). The motility of WT iNeutrophils was also impaired on the ARPC1B KO endothelium, suggesting that ARPC1B in endothelium regulates the motile behavior of iNeutrophils. The combined KO of ARPC1B in iNeutrophils and endothelium had the most dramatic effect on iNeutrophil migration (Fig. 4C). While ARPC1B deficiency in iNeutrophils induced the greatest defect in iNeutrophil motility, endothelial deficiency of ARPC1B was sufficient to induce a significant defect in iNeutrophil motility (Fig. 4D). Taken together, our findings show that ARPC1B in both neutrophils and endothelium regulate iNeutrophil interactions with endothelium.

**Figure 4.**
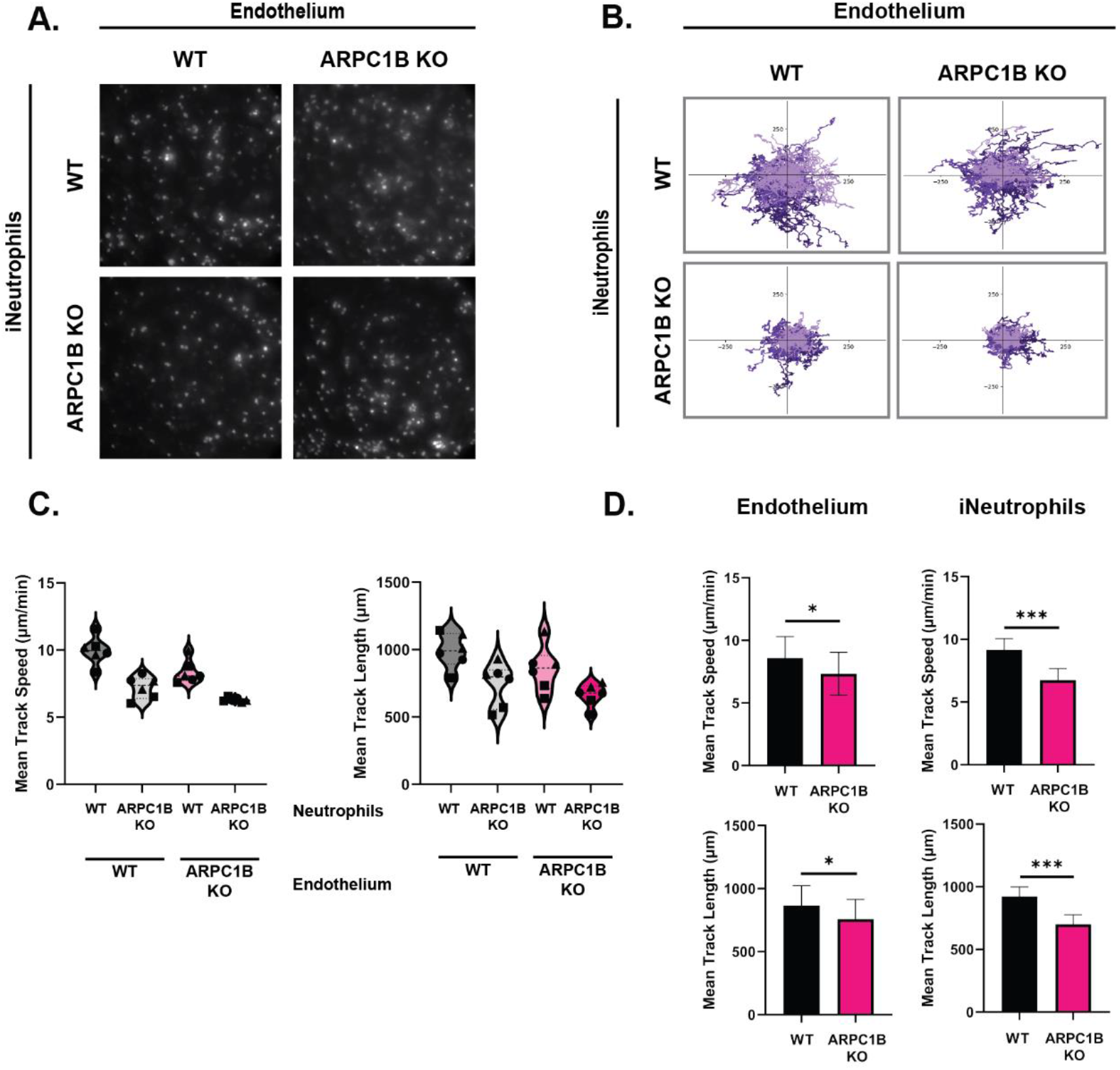
ARPC1B deficiency in both iNeutrophils and endothelium impairs neutrophil migration. (A) Representative images of iNeutrophil behavior after 2.5 hrs in contact with WT or ARPC1B-/-endothelium. (B) Compiled rose plots of all replicates 20-25 cells were tracked per condition and plotted, with color coding based on biological replicate. (C) Distribution of iNeutrophil velocity and total distance for all replicates. (D) Bar graphs represent the mean values of mean track speed and total distance traveled across conditions for endothelial cells with or without ARPC1B. *p* values were calculated by linear mixed effects model *, p < 0.05; **, p < 0.01; ***, p < 0.001; ****, p < 0.0001.

Here we show that ARPC1B in both neutrophils and endothelium drive the abnormal motility of neutrophils and their interactions with the endothelium. This study addresses a gap in understanding human neutrophil function because primary human neutrophils are terminally differentiated cells that are not genetically tractable. iNeutrophils offer the opportunity to genetically modify human neutrophils to understand neutrophil function and migration. Importantly, the motility and actin cytoskeletal defects found in patients with ARPC1B deficiency were more closely recapitulated in the iNeutrophils than in the commonly used HL-60 derived cancer cell line, PLB-985 cells.

There is substantial interest in understanding the dynamic formation of the leading edge during directed neutrophil migration. High speed imaging of migrating HL-60 cells has recently shown that Arp2/3-mediated actin branching optimizes lamellipodia shape to enable efficient cell migration (Garner & Theriot, 2022). Indeed, previous work from Weiner and colleagues (Servant et al., 1999) demonstrated that the Arp2/3 complex localizes to the leading edge where it colocalizes with sites of actin polymerization. ARPC1B is a component of this complex that is expressed specifically in hematopoietic cells. Previous studies have shown that ARPC1B is better at promoting actin assembly than other components of the ARP2/3 complex both in vitro and in cells (Abella et al., 2016). Here we show that ARPC1B-deficient iNeutrophils have impaired migration on 2 dimensions and that there is a switch from the formation of pseudopodia to the generation of elongated filopodia. These findings are consistent with the idea that ARPC1B is necessary for the rapid assembly of actin filaments at the leading edge necessary for the formation of pseudopodia in motile neutrophils. In addition to ARPC1B, a recent report showed that deficiency of the ARP2/3 component ARPC5 also leads to impaired neutrophil migration, further supporting a critical role for both ARPC1B and ARPC5 in neutrophil migration (Nunes-Santos et al., 2023).

Our findings also demonstrate that ARC1B in endothelial cells is sufficient to drive the abnormal interactions between neutrophils and the endothelium. We show that primary human neutrophils have impaired transmigration across ARPC1B-deficient endothelium in response to bacteria outside the lumen, even though the integrity of the lumen remains intact. This finding provides evidence that the immune deficiency and susceptibility to infection in these patients may also be mediated by abnormal endothelial regulation in the absence of ARPC1B. Indeed, we also found that WT iNeutrophils display reduced migration on KO endothelium and that this phenotype is further exacerbated if ARPC1B is absent in both the neutrophils and endothelium. The role of ARPC1B in endothelium may in part be mediated by the increased expression of inflammatory mediators in these cells. A recent report of a constitutively active Lyn kinase provides another example of endothelial cells functioning as drivers of neutrophilic vasculitis (de Jesus et al., 2023).

In summary, we have demonstrated that iNeutrophils provide a robust system to study neutrophil migration and model monogenic neutrophil inflammatory disorders. The findings support a key role for ARPC1B in regulating neutrophil actin cytoskeletal organization downstream of chemoattractant stimulation necessary for directed migration. The idea that endothelial ARPC1B can also drive neutrophil behavior and migration provides new insight into the underlying etiology of vasculitis syndromes.

## Materials and Methods

### Stem Cell Culture and Neutrophil Differentiation

Neutrophils were differentiated from bone marrow derived hiPSCs as previously described (Brok-Volchanskaya et al., 2019). Briefly, bone marrow-derived IISH2i-BM9 were obtained from WiCell (Madison, WI). hiPSCs were cultured on Matrigel-coated tissue culture plates in mTeSR™ Plus medium (STEMCELL Technologies, Vancouver, Canada).

To induce hemogenic endothelium, hiPSCs are transfected with ETV2 mmRNA in mTeSR™-E8™ complete media using TransIT reagent and mRNA boost ((Mirus Bio) One hour prior to transfection, cells were detached by TrypLE Select (LifeTech) to singularize. Cells were re-plated onto collagen (2.4ug/ml) (Type IV from human placenta Sigma Cat# C5533) coated plates in E8 with 10uM ROCK inhibitor (ROCKi; Tocris Y-27632) for transfection according to manufacturer’s protocol. One day following transfection, media was changed to StemLineII (Sigma Cat# S0192) media with 20ng/mL VEGF-165 and 10ng/mL FGF to induce differentiation into hemogenic endothelial cells. We refer to this media cocktail as Media A. After two days, the media was changed to differentiate the cells into common myeloid progenitors (CMPs) with StemLineII media supplemented with FGF2 (20 ng/mL), GM-CSF (25 ng/mL) (PeproTech), and UM171 (50 nM; Xcess Biosciences). We refer to this media cocktail as Media B. On days 8-10, floating cells were gently harvested and used for terminal neutrophil differentiation. These cells were cultured in StemSpan SFEM II medium (STEMCELL Technologies), supplemented with GlutaMAX 100X (Thermo Fisher Scientific), ExCyte 0.2% (Merck Millipore), human G-CSF (150 ng/mL; PeproTech), and Am580 retinoic acid agonist 2.5 μM (STEMCELL Technologies) at 2 × 10^6^ cells/mL density. We refer to this media cocktail as Media C. After 4 days, fresh Media C was added on the top of the cells. Neutrophils were harvested from the supernatant 8-10 days after plating in Media C.

## Generation of ARPC1B-/-PLB-985 cells

To generate PLB-985 Arpc1b CRISPR knockout line (Ran et al., 2013), oligos were designed to target the following sequence in exon 3: CTCGTGCACCTTGGTCCATTTGG

primer1: 5’ – CACCGCTCGTGCACCTTGGTCCATT – 3’

primer2: 5’ – AAACAATGGACCAAGGTGCACGAGC – 3’

Oligos were annealed and cloned into vector PX458 (SpCas9-2A-GFP) from Addgene #48138. pSpCas9(BB)-2A-GFP (PX458) was a gift from Feng Zhang (Addgene plasmid # 48138 ; http://n2t.net/addgene:48138 ; RRID:Addgene_48138). Vector was electroporated into PLBs using Nucleofector® II with solution V and program C-023. Cells were allowed to recover and then sorted for GFP-expressing population using cell sorter FACSAria (BD).

## Generation of ARPC1B-/-BM9iPSCs

To generate Arpc1b knockout BM9 iPSCs, two single guide RNAs were designed in the CRISPR design tool (Synthego) to target ARPC1B exon 3:

sgRNA1: 5’ – CTCATGGTTGTTGGGGCAGA – 3’

sgRNA2: 5’ – GACCAAGGTGCACGAGCTC – 3’.

Prior to nucelofection, BM9 iPSCs were treated with 10 µM ROCKi, detached by TrypLE Select (LifeTech), and singularized by pipetting. 2.5ug of each sgNRAs and 5ug of Cas9 protein (PNA Bio) were incubated together for 10 minutes, then the cells were nucleofected (protocol B16) using the Human Stem Cell Nucleofector Kit 2 (Lonza, #VPH-5022). Cells were plated at a low density on a Matrigel coated 10cm-plate at 25 cells/cm^2^ in mTeSR™ Plus media with 1xCloneR supplement (StemCell #05888). After 2 days, media was changed to mTeSR™ Plus media alone. Individual colonies were picked after 7 days and further expanded. To confirm biallelic mutation of the ARPC1B gene, genomic DNA was extracted from individual clones, then screened by PCR for the acquisition of a 66 bp deletion in the wild-type ARPC1B allele using sequencing primers primer1: 5’-CCTGCAAGGCAGAAGAGGAG-3’ and primer2: 5’-GGGCAGATGCTCCACAATG-3’. We selected a single clone (#4) for further experimentation.

### Human Neutrophil Isolation

Human blood was obtained from volunteering donors with informed written consent through a protocol that was approved by the Internal Review Board of the University of Wisconsin-Madison. Neutrophils were isolated using MACSxpress negative antibody selection kit and purified with the MACSxpress erythrocyte depletion kit (Miltenyi Biotec, Inc., Auburn, CA), following manufacturer’s instructions. Isolated neutrophils were resuspended in modified HBSS (+0.1% HSA +10mM HEPES) and utilized for various experiments. Incubations involving neutrophils were performed at 37°C with 5% CO2.

### Lumen protocol

The LumeNEXT devices were fabricated as previously described (Jiménez-Torres 2016). Briefly, the device was formed by patterning 2 polydimethylsiloxane (PDMS) (Dow) layers from SU-8 silicon masters (MicroChem) and bonding them using oxygen plasma (Diener Electronic Femto Plasma Surface System) onto a glass-bottom MatTek dish with a PDMS rod in the chamber (Supplemental Methods).

### iEC culture

iECs were generated using mmETV2 transfection through the hemogenic endothelium phase as described (see *Stem Cell Culture and Neutrophil Differentiation*). Briefly, bone marrow-derived IISH2i-BM9 cells were transfected with ETV2 mmRNA and cultured in media A for 1 day. Forty eight hours after transfection, media was changed to Vasculife Basal maintenance media (LifeLine Cell Technologies) supplemented with iCell-Endothelial Cells Medium Supplement (CDI). iECs were plated on cell culture-treated flasks preincubated with 30 µg/mL bovine fibronectin (Sigma-Aldrich F1141). iECs were passaged at 80% confluency and used through passage 5.

### Device and collagen preparation

LumeNEXT devices were prepared as previously described (Ingram 2018). Briefly, collagen I (Corning) was neutralized to pH 7.4 (3 mg/mL), pipetted into the devices and around PDMS rods. After polymerization, the PDMS rods were pulled from the chambers, leaving behind a lumen. Lumens were functionalized with 30 µg/mL bovine fibronectin (Sigma-Aldrich) and seeded with iECs at 2.5 × 10^4^ cells/µL. Lumens were flipped every 10 minutes in the incubator, and after 2 flips, unadhered cells were aspirated and replaced with fresh media. Lumens were cultured for 24 hours with media exchanges twice daily.

### Diffusion experiments

The media in the lumen was exchanged with 1:1 phosphate-buffered saline (PBS):iEC media. *P aeruginosa* or iEC media was added to the top port of the device and incubated for 2 hours and then set up for imaging. The PBS/iEC media was removed and replaced with a 50 µg/mL solution of 70 kDa fluorescein isothiocyanate (FITC)-Dextran (Sigma-Aldrich).

### Flow Cytometry

Flow cytometry analysis was used to evaluate expression of cell lineage markers on iPSC-derived neutrophils and endothelial cells. Cells were stained in PBS+ 1%HSA media +Brilliant Buffer (Thermo Fisher #00-4409-42) and Human TruStain FcX Fc Receptor Blocking Solution (Biolegend #422302), then fixed with 2% PFA. Data acquisition was performed on an Aurora Cytometer (CytekBio). Antibodies used in this study can be found in Supplementary Table 1. Forward and side-scatter parameters identified single cells and then live cells were identified with Ghost Dye Red 780 or Zombie NIR dye. Myeloid cells were identified by CD11b+ expression and neutrophils were identified by CD15+ or CD15+CD16+ expression. Monocytes were identified as CD14+ cells. Data were analyzed using FlowJo Software (v10.8.1). Endothelial cells were identified by CD31+, CD54+ and CD144+ expression.

### Chemotaxis

Chemotaxis was assessed using a microfluidic device as described previously (Berthier et al., 2010; Yamahashi et al., 2015). In brief, polydimethylsiloxane (PDMS) devices were plasma treated and adhered to glass coverslips. Devices were coated with 10 μg/mL fibrinogen (Sigma) in PBS for 30 min at 37°C, 5% CO2. The devices were blocked with 2% BSA-PBS for 30 min at 37°C, 5% CO2, and then washed twice with mHBSS. Cells were stained with Calcein AM (Molecular Probes) in PBS for 10 min at room temperature followed by resuspension in modified HBSS (+0.1% HSA +10mM Hepes). Cells were seeded at 2.5 × 10/mL to allow adherence for 30 min before addition of chemoattractant. Then, 3uL of 1 μM fMLP (Sigma) chemoattractant was loaded into the input port of the microfluidic device. Cells were imaged every 30 seconds for 45–90 min on a Nikon Eclipse TE300 inverted fluorescent microscope with a 10× objective and an automated stage using MetaMorph software (Molecular Devices). Automated cell tracking analysis was done using Imaris software to calculate mean velocity, straightness, and mean displacement.

### Endothelial Crawling

Endothelial cells were plated in a 24 well plate pre-coated with 30ug/ml of bovine plasma fibronectin (Thermo-Fisher) and cultured for a minimum of 48 hours, until cells were confluent. Four hours prior to experiment, media was replaced with RPMI supplemented with 2% FBS. IPSC-derived neutrophils were harvested and stained for 10 minutes with Calcein AM in PBS and resuspended at 2e6/mL in 2% FBS-RPMI. 50 uL of neutrophil suspension was added to each well. Cells were imaged every 1 minute for 2 hours on a Nikon Eclipse TE300 inverted fluorescent microscope with a 10× objective and an automated stage using MetaMorph software (Molecular Devices). Following the completion of imaging, cells were incubated for an additional hour and supernatants were centrifuged and frozen for later experimentation. Cell tracking analysis was done using the Manual Tracking functionality in ImageJ, and basic motility metrics and rose plots were calculated using Python.

### Immunofluorescent imaging

iNeutrophils were stimulated and prepared for immunofluorescent imaging as previously described (Klemm et al., 2021). Acid-washed 22-mm glass circle coverslips were coated with 10 µg/ml fibronectin for at least 1 h at 37°C and then blocked for 30 min with 2% BSA-PBS. Cells (3 × 10^5^) in 500 µl 0.5%HSA-RPMI were seeded per coverslip in a 24-well plate and allowed to rest for 30 min. fMLP was added for a final concentration of 100 nM fMLP and cells were allowed to adhere for 30 minutes at 37°C. Media was aspirated and fixation was performed as follows: 1 ml of 37°C preheated 4% paraformaldehyde in PEM buffer (80mM PIPES, pH6.8; 5mM EGTA, pH 8.0; 2mM MgCl2) for 15 min at RT. This was aspirated and 0.25% Triton X-100 in PEM buffer was added for 10 min at 37°C. Coverslips were washed three times with PBS, blocked in 5%BSA-PBS for 60 min at RT, washed, and incubated in Rhodamine-phalloidin (Thermo Fisher Cat#R415) overnight at 4°C. Coverslips were washed and incubated in secondary antibody solution for 60 min at RT. Coverslips were washed, counterstained with Hoechst 33342 (Thermo Fisher H3570; 1:500) for 5–10 min, washed with ddH2O, and mounted on Rite-On Frosted Slides (Fisher Scientific; 3050-002) with ProLong Gold Antifade Mountant (Invitrogen; P36930). Slides were imaged on an upright Zeiss LSM 800 Laser Scanning Confocal Microscope equipped with a motorized stage and Airyscan module. Images were acquired with a 60× oil, NA 1.40 objective using the Airyscan acquisition mode. Images were processed in ZenBlue software (Zeiss) using the Airyscan processing method and maximum intensity projections generated for further analysis. Filopodia were quantified using the FiloQuant plugin in ImageJ (Jacquemet et al., 2017). The following parameters were used to identify average number and length of filopodia per cell:

### Real Time qPCR

iPSC-differentiated neutrophils were stimulated with LPS to evaluate expression of inflammatory cytokines. A 6 well TC treated plate was pre-coated with 10ug/ml fibronectin. Approximately 6 million cells resuspended in modified HBSS (mHBSS) were plated per well. Cells were allowed to rest for 30min at 37°C before stimulation. Final concentration of 200ng/mL E. coli LPS (Sigma # L2755) was added to appropriate wells and the plate was incubated for 2 hours at 37°C. Floating cells were collected and spun down at 300xg. To collect RNA, 1ml of Trizol (Invitrogen) was added to adherent cells and then combined with the pelleted floating cell sample. Samples were pipetted to mix and then stored at -80°C until RNA extraction. RNA was isolated by Trizol extraction following manufacturer’s instructions (Invitrogen #15596026). Collected RNA was stored at -80°C until cDNA preparation. cDNA was synthesized using oligo-dT primers and the Superscript III First Strand Synthesis kit (Invitrogen #18080-051) following manufacturer’s instructions. cDNA was used as the template for quantitative PCR (qPCR) using FastStart Essential Green DNA Master (Roche) and a LightCycler96 (Roche). Data were normalized to ef1a within each sample using the ΔΔCq method (48). Fold-change represents the change in cytokine expression over the unstimulated WT sample. All qPCR primers are listed in Supplementary Table 2.

### Statistical Analysis

All experiments and statistical analyses represent at least three independent biological replicates (N). Prior to statistical analysis, all datasets were analyzed for Normality using the Kolmogorov-Smirnov test. All chemotaxis experiments, qPCR gene expression, and flow cytometry were analyzed using unpaired student t-test or Kolmogorov-Smirnov test, depending on whether the dataset passed Normality. The above analyses were conducted using GraphPad Prism (v9). Endothelial crawling experiments, filopodia analysis, and integrated density of F-actin were analyzed using a linear mixed effects model to account for clustering of measurements within the same day, and p-values were reported for effect sizes between endothelial cells and neutrophils. Linear mixed effects models were constructed and tested using R (v4.3.0; R Core Team 2023). All graphical representations of data were created in GraphPad Prism (v9) and figures were ultimately assembled using Adobe Illustrator (Adobe version 23.0.6).

## Acknowledgements

Research reported in this publication was supported by the National Institute of General Medical Sciences (NIGMS) of the National Institutes of Health under Award Number R35 GM118027 to Anna Huttenlocher. The content is solely the responsibility of the authors and does not necessarily represent the official views of the NIH. The funders had no role in study design, data collection and analysis, decision to publish, or preparation of the manuscript. The content is solely the responsibility of the authors and does not necessarily represent the official views of the National Institutes of Health.

## Author Contributions

A.N.P, D.A.B., and J.C. performed experiments. A.N.P, D.A.B., M.L., D.J.B., and A.H analyzed and interpreted data. A.N.P. and A.H planned and designed the research and wrote the paper.

## Correspondence

Anna Huttenlocher, University of Wisconsin–Madison, 1550 Linden Dr, Room 4225, 411 Madison, WI 53706;

