## Supplemental Data for "Neutrophil motility is regulated by both cell intrinsic and endothelial cell ARPC1B"

### Supplemental figure

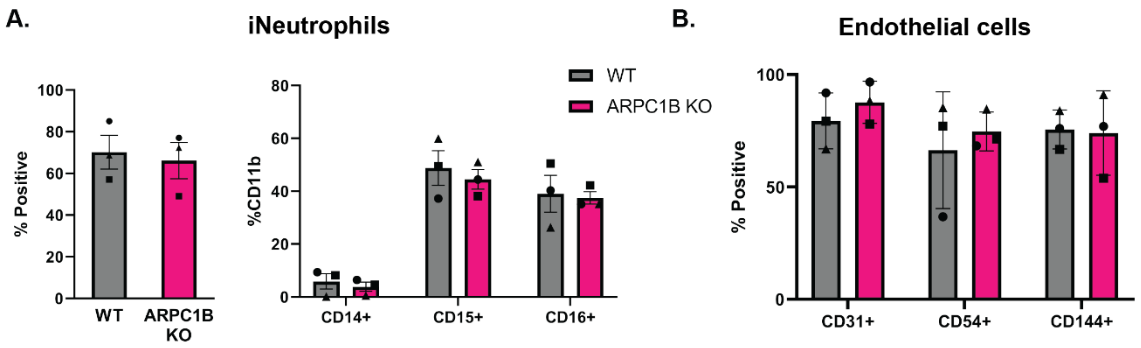


**Supplemental Figure 1. Flow cytometry staining of WT and ARPC1B KO iNeutrophils and endothelium**

(A) Flow cytometry staining of differentiated neutrophils. Cells were first gated on live singlets and CD11b+ expression was quantified. CD14, CD15, and CD16 expression is quantified as a percentage of CD11b+ population. (B) Flow cytometry staining of hemogenic endothelium obtained during differentiation process. Cells were first gated on live singlets.

### Movies

#### Movie 1. PLB-985 cells

Representative time-lapse imaging of WT vs ARPC1B KO PLB-985 cells, corresponding to Figure 1. Cells are migrating on 10 μg/mL fibronectin towards an fMLP gradient (1 μM) in a microfluidic device, originating at the bottom of the frames. The frame rate is 10 frames per second. Images were acquired every 30 seconds for 45 minutes and movies were trimmed to start 15 minutes after the chemoattractant was loaded. Cells were labeled with Calcein AM. Scale bar is 100 μm.

**Movie 2. iNeutrophil motility to fMLP gradient**

Representative time-lapse imaging of WT vs ARPC1B KO iNeutrophils, corresponding to Figure 2. Cells are migrating on 10 μg/mL towards an fMLP gradient (1 μM) in a microfluidic device, originating at the bottom of the frames. The frame rate is 10 frames per second. Images were acquired every 30 seconds for 45 minutes and movies were trimmed to start 15 minutes after the chemoattractant was loaded. Cells were labeled with Calcein AM. Scale bar is 100 μm.

**Movie 3. Primary human neutrophil motility out of modified IPS-derived endothelial lumen**

Representative time-lapse imaging of human primary neutrophils out of a WT or ARPC1B KO IPS-derived endothelial lumen using the LumeNext system. Neutrophils are migrating to a source of *P. aeruginosa* at the top of the devices. Images were acquired every 10 minutes for 16 hours. Neutrophils were labeled with Calcein AM.

**Supplementary Table 1: Flow cytometry antibodies used**

| Target | Fluor | Clone | Cat# | Vendor |
|---|---|---|---|---|
| CD11b | PE | HI10a | 312203 | Biolegend |
| CD14 | BUV805 | M5E2 | 612902 | BD Biosciences |
| CD15 | APC | W6D3 | 323008 | Biolegend |
| CD16 | BV711 | 3G8 | 563127 | BD Biosciences |
| CD31 | PE | WM59 | 303105 | Biolegend |
| CD54 | AF700 | 353125 | 353125 | Biolegend |
| CD144 | APC | BV9 | 348507 | Biolegend |
| Ghost Dye Red | 780 | | 13-0865-T100 | Tonbio Biosciences |
| Zombie NIR | 746 | | 423105 | FisherScientific |
| Brilliant Buffer | | | 00-4409-42 | Thermo Fisher |

| | | | | |
|---|---|---|---|---|
| Human TruStain FcX Fc Receptor Blocking Solution | | | 422302 | Biolegend |
| Ultracomp ebeads | | | 501129040 | Thermo Fisher |

**Supplementary Table 2: qPCR primers used**

| Gene | Sequence | Source |
|---|---|---|
| EF1α | F - TGGTATTGGTACTGTTCCTG | KiCqStart SYBR Green Primers |
| | R - CTTCACTCAAAGCTTCATGG | |
| IL1β | F - CTAAACAGATGAAGTGCTCC | KiCqStart SYBR Green Primers |
| | R - GGTCATTCTCCTGGAAGG | |
| IL6 | F - TCTCCACAAGCGCCTTCG | |
| | R - CTCAGGGCTGAGATGCCG | |
| IL8 | F - TGTAAACATGACTTCCAAGC | KiCqStart SYBR Green Primers |
| | R - AAAACTGCACCTTCACAC | |
| TNFα | F - CCTCTCTCTAATCAGCCCTCTG | PrimerDB |
| | R - GAGGACCTGGGAGTAGATGAG | |
| CDH5/VE-cadherin | F - TTGGAACCAGATGCACATTGAT | (Wang et al., 2012) |
| | R - TCTTGCGACTCACGCTTGAC | |
| ICAM-1 | F - ATGCCCAGACATCTGTGTCC | (Wang et al., 2012) |
| | R - GGGGTCTCTATGCCCAACAA | |
